# Measuring Mechanodynamics using an Unsupported Epithelial Monolayer Grown at an Air-Water Interface

**DOI:** 10.1101/052555

**Authors:** Corinne Gullekson, Matthew Walker, James L. Harden, Andrew E. Pelling

## Abstract

Actomyosin contraction and relaxation in a monolayer is a fundamental biophysical process in development and homeostasis. Current methods used to characterize the mechanodynamics of monolayers often involve cells grown on solid supports such as glass or gels. The results of these studies are fundamentally influenced by these supporting structures. Here, we describe a new methodology for measuring the mechanodynamics of epithelial monolayers by culturing cells at an air-liquid interface. These model monolayers are grown in the absence of any supporting structures removing cell-substrate effects. This method’s potential was evaluated by observing and quantifying the generation and release of internal stresses upon actomyosin contraction (320±50Pa) and relaxation (190±40Pa) in response to chemical treatments. Although unsupported monolayers exhibited clear major and minor strain axes, they were not correlated to nuclear alignment as observed when the monolayers were grown on soft deformable gels. It was also observed that both gels and glass substrates led to the promotion of long-range cell nuclei alignment not seen in the hanging drop model. This new approach provides us with a picture of basal actomyosin mechanodynamics in a simplified system allowing us to infer how the presence of a substrate impacts contractility and long-range multi-cellular organization and dynamics.

## INTRODUCTION

Contractility is involved in the remodeling and organization of the cell interior and plays a major role in multicellular morphogenesis over long distances and timescales (Rauzi *et al.*, 2010; Harris *et al.*, 2012; Roh-Johnson and Shemer, 2012). There is increasing interest in understanding how contractility manifests in multi-cellular systems (Nelson *et al.*, 2005; Fernandez-Gonzalez *et al.*, 2009; Rauzi *et al.*, 2010). Contractile forces generated by the actomyosin cytoskeleton within individual cells collectively generate tissue-level forces (Vicente-Manzanares *et al.*, 2009; Martin *et al.*, 2010). A major challenge in developmental biology is to understand how cytoskeletal activity is orchestrated to produce higher order tissue organization (Fernandez-Gonzalez *et al.*, 2009). Therefore, understanding tissue morphogenesis requires determining how cellular forces are integrated across cells and tissues (Martin *et al.*, 2010).

Actomyosin contraction is one of the major sources of internal force inside of the cell. The actin cytoskeleton in epithelial monolayers is highly dependant on cell attachment. In epithelia, the cell-cell attachment points, adherens junctions, link a thick circumferential ring of actin and myosin around each cell, aligned with the cell borders (Owaribe *et al.*, 1981; Yonemura *et al.*, 1995). The contractility of these marginal actin bundles is used for morphogenesis facilitating epithelial sheet bending and invagination (Leptin, 2005; Lecuit and Lenne, 2007). The substrate attachment points, focal adhesions, link actin stress fibers. Cell contractility is commonly described and investigated in terms of changes in cortical cell elasticity as well as traction dynamics, the resistance of the substrate to deformation, on flexible substrates (Discher *et al.*, 2005; Rauzi *et al.*, 2008). Generally changes in cortical cell elasticity and traction dynamics are linked to sensing of the mechanical microenvironment (Vogel and Sheetz, 2006; Trichet *et al.*, 2012). Traction dynamics are linked to cortical elasticity, focal adhesion organization and cell morphology (Pelham and Wang, 1997; Discher *et al.*, 2005; Solon *et al.*, 2007). Often fundamental studies of actomyosin contraction and relaxation are examined in situations where cells are grown on, or embedded in, flexible substrates. (Discher *et al.*, 2005; Ghibaudo *et al.*, 2009; Huh *et al.*, 2010; Tremblay *et al.*, 2013). Currently there is intense interest in understanding the interplay and feedback between the mechanical properties of such substrates and contractility (Trichet *et al.*, 2012). Although such studies typically employ substrates with systematically altered mechanical properties, the substrate is always present. Therefore, it remains difficult to assess the intrinsic internal mechanical dynamics and properties of cells in the absence of a mechanically supporting surface.

There has been recent interest in intrinsic mechanical properties of substrate-free epithelial sheets (Harris *et al.*, 2012, 2013). Measurements of the elasticity and failure of monolayers suspended between two flexible rods provided valuable insights into the understanding of epithelial mechanics (Harris *et al.*, 2012, 2013). Importantly, the measurements yielded mechanical properties of the near substrate-free cell sheet. With tensile testing of the suspended cultured monolayers, it was determined that rupture of intercellular junctions occurs after doubling the monolayer in length with an average force approximately nine times larger than measured in pairs of isolated cells (Chu *et al.*, 2004; Harris *et al.*, 2012), demonstrating that epithelial organization leads to a strong enhancement in the integrity of the tissue. However this suspended monolayer is still mechanically constrained at suspension points that provide an external form of confinement and pre-stress.

This begs the question, how are cell contractility, the organization of the cell interior and long range structural remodeling altered in the absence of mechanically confining or supporting structures in a multi-cellular system? Therefore, the objective of this study was to develop a new methodology to characterize actomyosin mechanodynamics in a multi-cellular system when no resistance is offered by the surroundings. Understanding these dynamics could provide us with a picture of basal actomyosin mechanodynamics in a simplified model multi-cellular system. This method will also allow us to indirectly infer how the presence of a substrate impacts actomyosin contractility and long-range multi-cellular organization and dynamics.

Here, we employed an in-vitro hanging drop culture protocol. We created Madine-Darby Canine Kidney (MDCK) epithelial monolayer clusters at an air-water interface. Employing laser scanning confocal microscopy, cell nuclei within the monolayer were used as fiduciary markers to determine the internal strain developed in the clusters during basal and chemically-induced contractile changes. We validated our method using two alternate methods for tracking cell deformation: 1) tracking the cell membrane boundary of the cluster and 2) tracking cell boundaries marked with GFP-actin. Importantly, this new methodology for studying the mechanodynamics of unsupported epithelial sheets opens the door for several future research directions including investigations of the apico-basal polarization, myosin distribution and actin ordering in the unsupported monolayer. This approach allows one to observe the results of forces that are important for many processes that take place in development (Heisenberg and Bellaïche, 2013) and may advance the understanding of how those forces are being transmitted to neighbouring cells, and how they are integrated to trigger global changes in tissue shape. The results of this study might also be relevant to 3D cell culture where hard surfaces are not present.

## RESULTS

### Deformation of Monolayer Clusters on Solid Substrates

To examine the influence of a mechanically supporting substrate, monolayer clusters were created on hard glass substrates and soft polyacrylamide (PA) gels (4.8 kPa Young’s modulus). These clusters naturally occur in MDCK cell culture at low confluence. The clusters on glass were fixed and stained for DNA, actin and vinculin (Figure 1A). A strong actin signal was localized in the perimeter of the cluster. Similarly the vinculin also localized around the cluster perimeter. When epithelial cells form a cluster on solid substrates, more cellular adhesions and force generation appear at the cluster perimeter (Notbohm *et al.*, 2012; Mertz *et al.*, 2013; Ng *et al.*, 2014). In order to induce cluster relaxation and contraction in the clusters we employed the agents blebbistatin and CalA. Blebbistatin is a myosin II inhibitor that inhibits contraction and disrupts contractile filament organization (Lemmon *et al.*, 2009). CalA is a phosphatase inhibitor that is well known to induce contraction (Fernandez-Gonzalez *et al.*, 2009; Lemmon *et al.*, 2009). Clusters on glass were live stained with Hoechst and the nuclei were tracked with the addition of drugs in order to determine strain in the cluster (Figure 2), as described in the Materials and Methods. There was no significant difference in the mean strains of the clusters on glass with the different drug treatments (p>0.12) (Figure 1B). To investigate the deformation on a softer substrate, clusters were cultured on PA gels. Consistent with the glass substrates, there was no significant difference in the mean strains of the clusters on the gels with the different drug treatments (p>0.07) (Figure 1C). To more thoroughly examine the intrinsic deformation of a contracting or relaxing monolayer the solid substrate should be removed, for this reason the hanging drop method was introduced.

**Figure 1.**
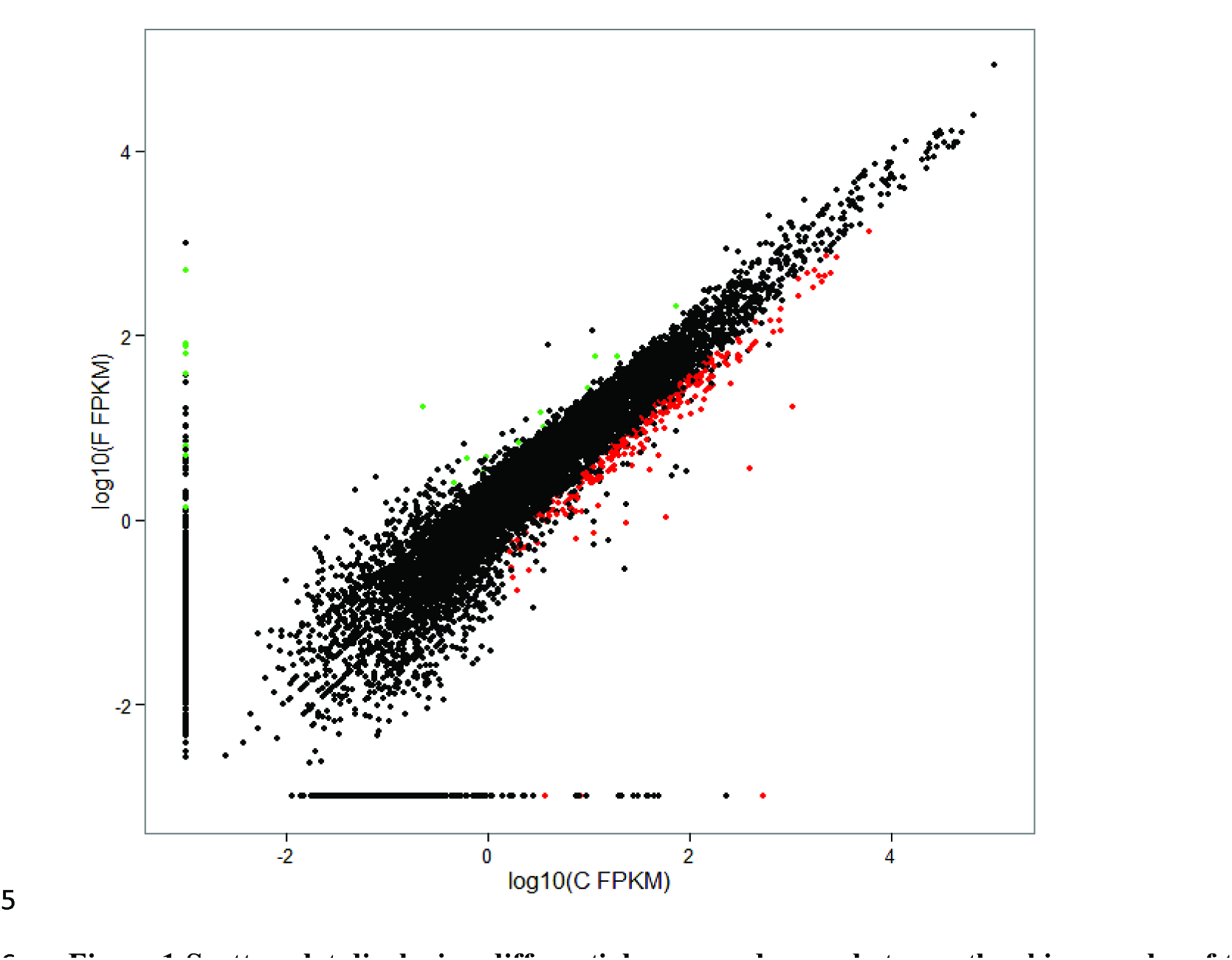
Deformation of the cluster on glass and PA. (A) Confocal images of the actin (left), vinculin (middle) and merged (right, actin=red, vinculin=green, nucleus=blue) of a cluster of MDCK cells on glass. (B) Major E_1_ minor E_2_ and mean E_M_ strains of cell clusters on glass after the addition of blebbistatin (red), media (blue) or calyliculin A (green). n=11, 9 and 12, no significance. (C) Major E_1_ minor E_2_ and mean E_M_ strains of cell clusters on a PA gel after the addition of blebbistatin (red), media (blue) or calyliculin A (green). n=15, 10 and 15, no significance. Bar, 10 μm.

**Figure 2.**
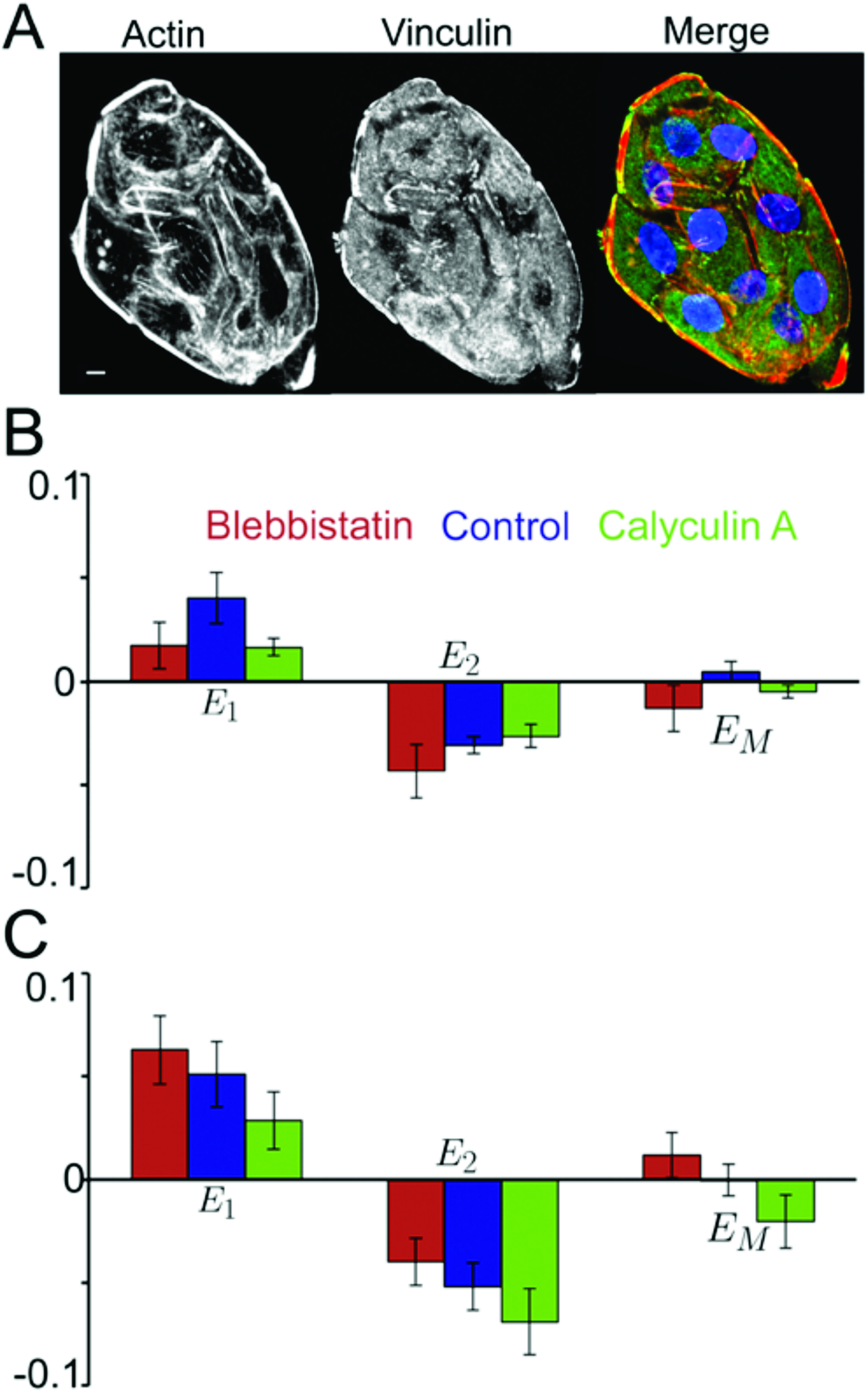
The process used to analyze the images. (A) Maximum intensity Z-projections of the before and after confocal images of cells in a hanging drop used for a control. (B) The positions of the nuclei in the before (blue) and after (red) images determined by threshholding. The raw nuclear positions (left) are shown as well as the positions after the translation (middle) and after the rotation (right) of the after positions. (C) Positions of the before (blue) and after (red) nuclei in a mesh over a mean strain map of the cluster. The major and minor principle axes (E_1_ and E_2_) of the cluster are in the y and x directions respectively. Maps are shown for the control (left) case as well as CalA (middle) and blebbistatin (right) treatments. Bar, 10 μm.

### Formation of a cell monolayer cluster

As MDCK cells migrate and proliferate in 2D culture, the cells form islands that eventually merge to form a continuous monolayer (Zegers *et al.*, 2003). In 3D culture, individual MDCK cells plated within an extracellular matrix gel will assemble into a hollow sphere that is lined by a monolayer of polarized epithelial cells (Zegers *et al.*, 2003). However, after incubation in a hanging drop, MDCK cells will form monolayer clusters of cells (Tchao, 1989; Eckhart *et al.*, 2003; Inoue *et al.*, 2005). This is in contrast to mesenchymal stem cells, which gradually coalesce into a single central spheroid along the lower surface of the drop (Bartosh, 2010). It has been shown that MDCK cells in a hanging drop will form a basement membrane-like sheet of cell-secreted proteins providing the matrix for the proliferating cells (Eckhart *et al.*, 2003; Inoue *et al.*, 2005). Previous immunohistochemical studies demonstrated the presence of type IV collagen, laminin 1, and laminin 5 in the membrane, and enzyme digestion experiments indicated that this membrane was sensitive to collagenase (Eckhart *et al.*, 2003). In previous studies, it has been shown that the tension carried by basement membranes are modest compared with the tension carried by the cellular portion of the epithelium (Wiebe *et al.*, 2005) and that the basement membrane is thinner and softer than the cellular component with a Young’s modulus of 7.5kPa (Last *et al.*, 2009) compared with 20kPa (Harris *et al.*, 2012). Moreover, this basement membrane is relatively free to move along the air-water interface. Therefore, this basement membrane-like sheet allows cells to form healthy monolayers while still being relatively unrestrictive to collective cellular deformation.

Epithelial monolayer clusters were grown in these hanging drops and their nuclei were used to track deformations during changes in contractility. The clusters were similar in number of cells (average of 23±2 cells per cluster) to the ones used in the solid substrate experiments (average of 21±2 cells per cluster). To label cell nuclei, we added the common nuclear stain, Hoechst 33342, to the drop with a pipette in the lid’s inverted state (Figure 3, A and B). In all cases, 1μL of a stock solution of blebbistatin, CalA or media (control) was added to the hanging drop for 30 minutes in a cell culture incubator. The nuclei of suspended clusters were imaged before and after the addition of the drugs or control. On average, only five monolayer clusters formed in each hanging drop and it was straightforward to image the same cluster before and after the 1μl addition.

**Figure 3.**
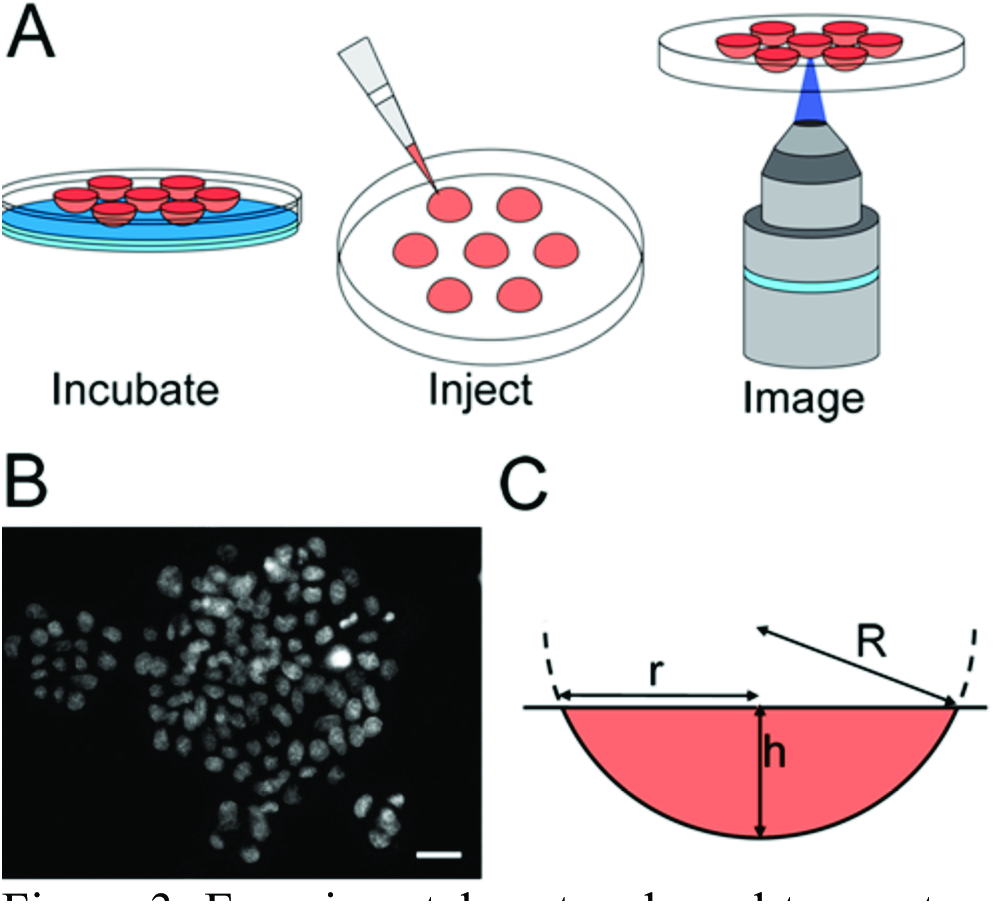
Experimental protocol used to create monolayer clusters in hanging drops. A) The process used to grow, stain and image the monolayer clusters before and after the addition of cytoskeletal drugs. 40μ1 droplets were incubated overnight with PBS in the bottom of the dish. The dish lid was flipped and 1 μ1 of Hoechst in media was added with a pipette. The droplets were then imaged with a confocal microscope. After this, drugs were added to the droplets, which were incubated for 30minutes and imaged again. B) A typical set of clusters found in a droplet after the addition of Hoechst. C) A diagram describing the shape of the droplet. The droplet shape can be approximated as a spherical cap with a radius of curvature, *R*. The radius at the droplet’s top, r, is 2.9±0.1mm. The approximate height of the droplet, *h*, is calculated with the known volume of the droplet, 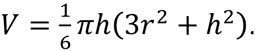. Bar, 20 μm.

### Hanging drop shape

The shape of the droplet observed in this set up can be approximated as a spherical cap (Figure 3C). The radius, r, at the droplet’s top, where it is attached to the dish lid, was measured optically at 2.9±0.1mm (n=7). The droplet volumes were measured using the mass of the droplets to be 39.49±0.04, 40.29±0.06 (after Hoechst addition) and 41.18±0.07μl (after drug addition, n=10). Using the spherical cap model, the height, h, and radius of curvature, R, of the initial droplet were calculated to be 2.5±0.1mm and 2.9±0.1mm respectively. With the addition of 2μl, the height of the droplet increased by 74.4±0.5μm and the radius of curvature decreased by 14±8μm (0.5% decrease). The average diameter of clusters was approximately 150μm. With the comparatively large radius of curvature, the center and edge of the cluster had a height difference of only 0.96±0.04μm. For this reason it was reasonable to approximate the curved bottom of the drop as a flat surface. Due to the small size of the droplet, scans were acquired quickly to avoid evaporation effects. Under the imaging power employed in this study, some evaporation of the droplet did occur, resulting in a height decrease of 0.24±0.02μm per second, which corresponds to a change in volume of 0.32±0.02μl per minute (Figure S1). For this reason, image collection was always maintained under 45 seconds. The radius of curvature only decreased by a maximum of 1.5±0.1μm during imaging, having little effect on the monolayer clusters’ curvature (under 0.05%).

### Deformation of Monolayer Clusters in Hanging Drops

The cytoskeletal drugs, blebbistatin and CalA, were used to relax and contract the cell clusters. The displacements of the cell nuclei were fit to determine the major and minor principle strains (E_1_ and E_2_) of the cluster. As a control, 1μl of media was added to the drops followed by incubation for 30 minutes between imaging. We did observe slight movement of nuclei within the cluster. However, the mean strain of the clusters in the control study was not significantly different than zero (p>0.89) (Figure 4). As expected there was no significant dilation or contraction. The myosin II inhibitor, blebbistatin, caused clusters to dilate appreciably. The mean strain was significantly higher (p<0.00001) than the control. Clusters also contracted with the addition of CalA resulting in a mean strain that was significantly lower (p<0.0004) than the control (Fig 4).

**Figure 4.**
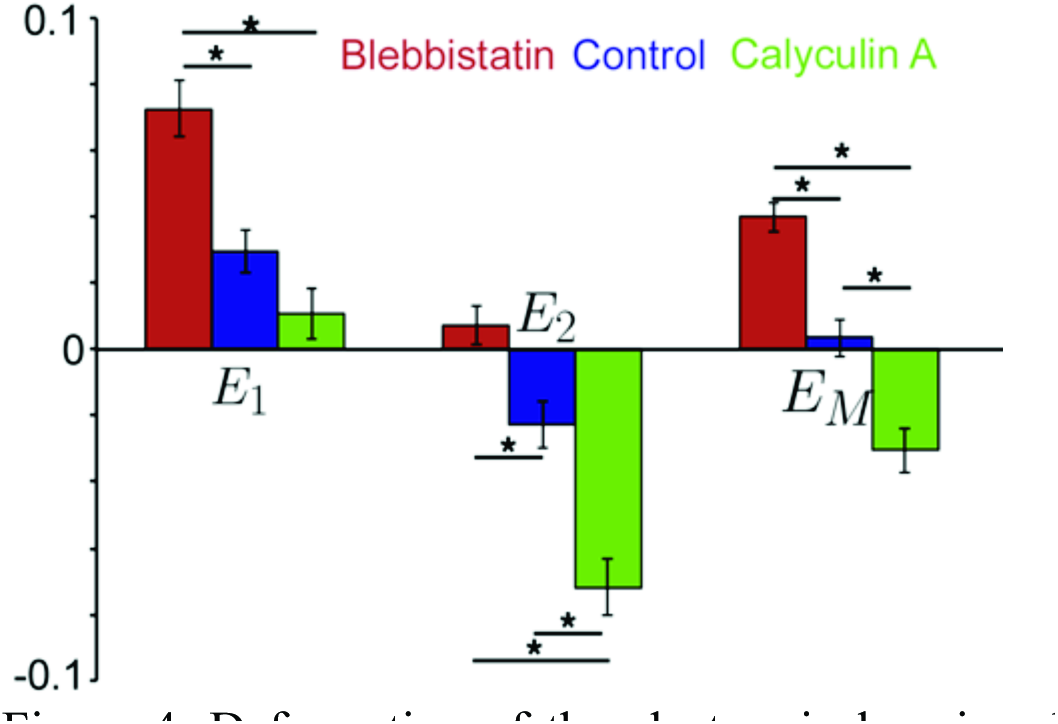
Deformation of the clusters in hanging drops with cytoskeletal drugs. Major (E_1_), minor (E_2_) and mean (E_M_) strains of cell clusters after the addition of blebbistatin (Red), media (blue) or CalA (green). n=22, p<0.05.

### Cluster Outline Deformation

To determine if the perimeter of the monolayer cluster itself deforms in a manner consistent with the changes in nuclei position, the cluster outline was tracked by fluorescently labeling cell membranes with WGA. Tracking points on the cluster perimeter allowed us to determine monolayer strain (Figure S2A). The major and minor strains in the cluster were calculated from nuclei positions as well as from its outline. The strains calculated using cell nuclei and cluster outlines were not significantly different for the untreated control (p>0.90), blebbistatin (p>0.85) and CalA treatments (p>0.82) (Figure S2B). This suggests the contraction and relaxation observed in the nuclear positions is a good indicator of total cluster deformation.

### Individual Cellular Strains

To determine individual cellular strains during cluster contraction and relaxation, cells transiently expressing actin-GFP were used to form suspended clusters. It was found that when a high percentage of cells were expressing actin-GFP, CalA treatments had no effect on contraction. Therefore, we adjusted the transfection efficiency to ∼25% of cells in order to create suspended clusters that exhibited the same mechanical dynamics as non-transfected cells. In this case, we felt confident that we would be able to examine individual cellular strains, albeit only on a sub-population of cells within each cluster. Distinct actin stress fibers were not evident but actin was found localized to the cell margins allowing us to easily determine cell boundaries (Figure 5, A and B). Experiments with blebbistatin, CalA and control conditions were performed once again and nuclei positions were tracked, as well as the deformation of single cells within the cluster.

**Figure 5.**
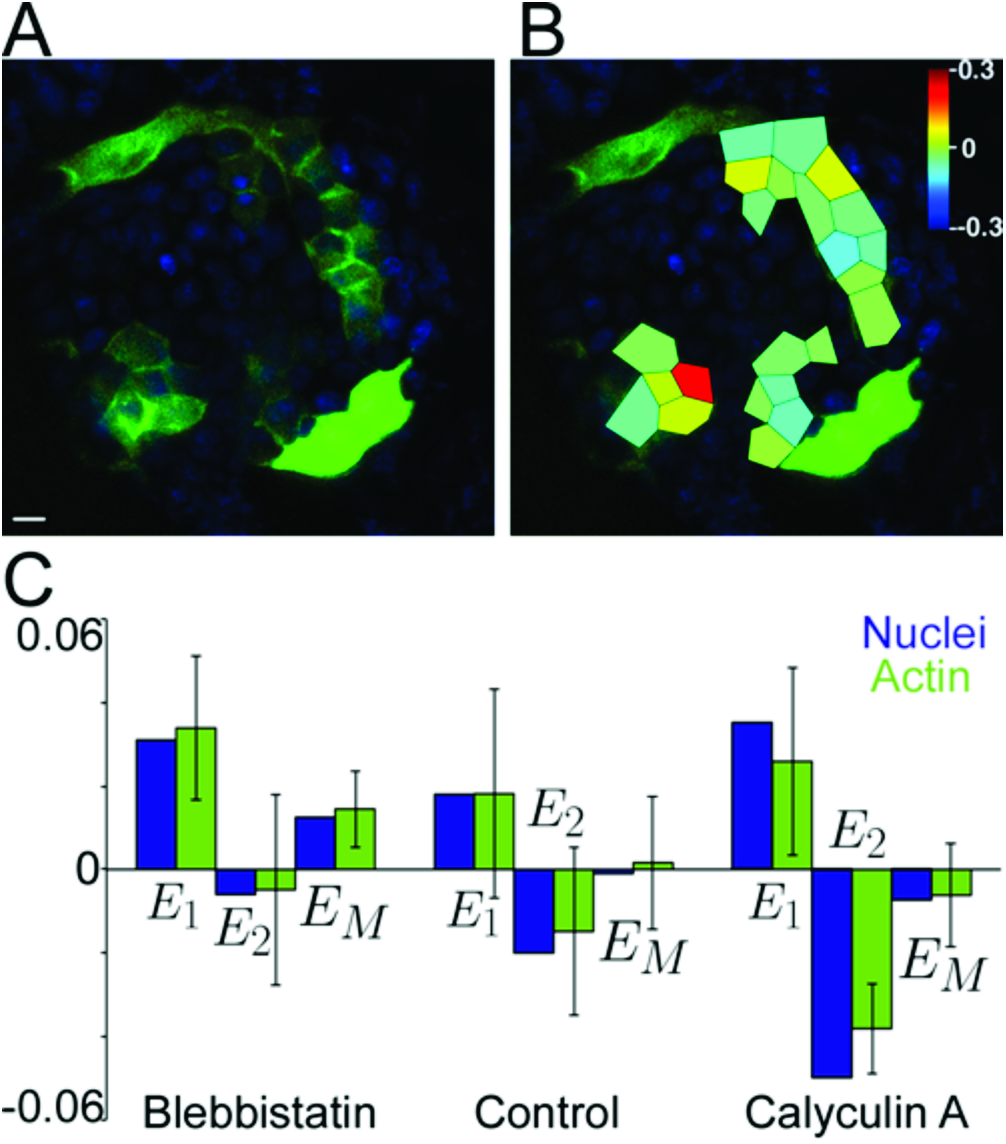
Cellular deformations inside a deforming hanging drop cluster. (A) The before image of a MDCK cluster stained with Hoechst (blue) partially transduced with GFP-actin (green). (B) The mean strain of each cell overlaid on the before image. (C) Major, minor and mean strain of cell clusters after the addition of blebbistatin, media, or CalA (blue) compared to the strain in the individual cells as calculated with the actin cell boundaries (green). Bar, 10 μm.

In the control, the cluster underwent normal basal remodelling as seen by nuclear displacements. With the addition of CalA and blebbistatin the clusters the nuclear displacements showed contraction and relaxation of the cluster as expected (Figure 5C). The mean of the strains in the individual cells were not significantly different than the cluster strains calculated by nuclear displacements in the control (p>0.97), blebbistatin (p>0.96) and the CalA treatments (p>0.98). This demonstrates that the monolayer cluster strains determined by nuclei correspond well to the deformation of the individual cells themselves.

### Substrate Effects on Strain

Comparing the blebbistatin or CalA induced strains on the glass or PA substrates to the hanging drop (Figure 6A) reveals that they were significantly muted (p<0.002 in both cases on glass, and p<0.04 for blebbistatin on PA) by the substrate. Interestingly however, there was no significant difference in the controls (p>0.87).

**Figure 6.**
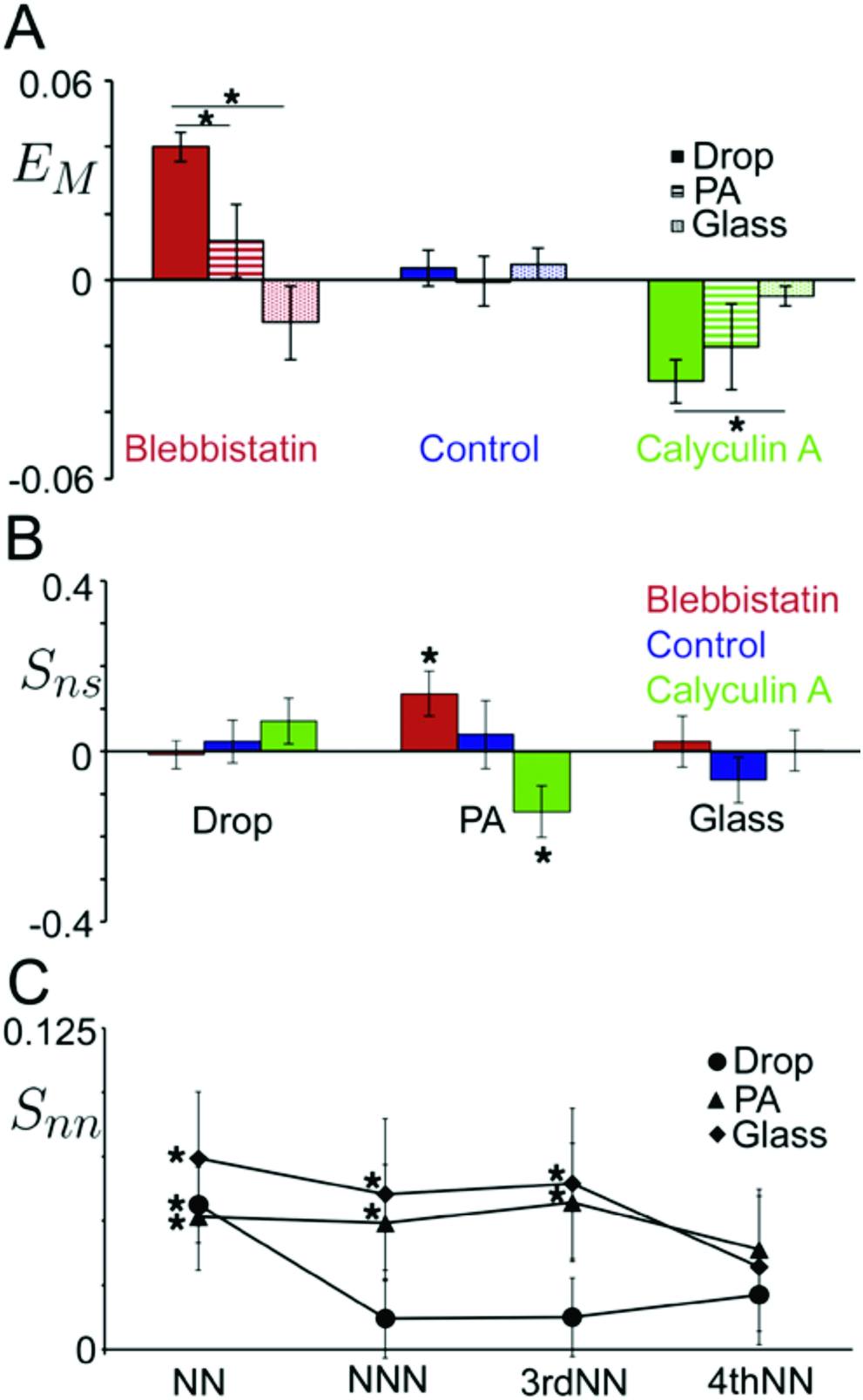
Substrate dependence on nuclear deformation (A) Mean E_M_ strains of cell clusters in the hanging drops (solid, n=22), on PA gels (stripes, n=15, 10 and 15) and glass (dots, n=11, 9 and 12) after the addition of blebbistatin (red), media (blue) or CalA (green). (B) Order parameter, *S*_*ns*_, for nuclei (defined by the angle between the major strain axis and the nuclear major axis) for blebbistatin (red), control (blue) and CalA (green) treatments for hanging drops, PA and glass. (C) Order parameters, *S*_*nn*_ for nuclei (defined by the angle between the nuclear major axis and its neighbor’s nuclear major axis) in hanging drops (circles), on PA (triangles) and glass (diamonds). The results are shown for nearest, next nearest, third nearest and fourth nearest neighbors. p< 0.05.

### Feature Orientation

In order to determine if the deformation of the cluster exhibited a dependence on structural features of the cells, nematic order parameters were calculated as described in the Materials and Methods. The order parameter for nuclear alignment with strain (*S*_*ns*_) was calculated for all the clusters (Figure 6B). The order parameters in the hanging drops were not significantly different than 0 (p>0.82, 0.64 and 0.20 for blebbistatin, control and CalA, respectively), suggesting no preferential alignment. On glass, the order parameters were also not significantly different than 0 (p>0.71, 0.25 and 0.97 for blebbistatin, control and CalA), respectively, suggesting no preferential alignment. However, since the strains were small on glass, the major strain axis was likely determined by basal remodeling of the cells and not the contraction or relaxation of the actomyosin cytoskeleton. On PA, the control order parameter was not significantly different than 0 (p>0.62), however the order parameters of treated clusters’ nuclei were significantly different than 0. In the blebbistatin treated clusters, the nuclei preferentially aligned with the major strain axis (p<0.04). In the CalA treated clusters, the nuclei preferentially aligned with the minor strain axis (p<0.04). Therefore, the nuclei tended to be aligned with the relaxation and contraction axes. This demonstrated that there was a correlation between nuclear alignment and cluster strain on PA gels that was not observed in hanging drops or on rigid glass substrates.

The order parameter *S*_*ls*_, calculated with the angle between the major length axis and the major strain axis of the cluster (Figure S3A), was not significantly greater than 0 for any treatment or substrate (p>0.09 in all cases). Another order parameter (*S*_*nl*_) examines if the nuclei were aligned with the major length axis on the cluster. Again, this parameter was not significantly greater than 0 for any treatment or substrate (p>0.57 in all cases) (Figure S3B). The average eccentricities of the nuclei grown on the three different substrates were not significantly different (p>0.15). The clusters also had varying major/minor length aspect ratios, however the mean strains observed in the monolayer clusters did not display any correlation with aspect ratio (Figure S4).

To examine the alignment of cells with their neighbors, the order parameters between nuclei (*S*_*nn*_) were calculated (Figure 5D). For all substrates, the order parameters for the nearest neighbors were significantly greater than 0 (p<0.002, 0.02 and 0.005 for hanging drops, PA and glass, respectively) suggesting a local alignment of cells in direct contact. For next nearest neighbors and third nearest neighbors, the order parameter was not significantly greater than 0 for hanging drops (p>0.4, 0.4), but was significantly greater than 0 for PA (p<0.03, 0.02) and glass (p<0.05, 0.03). After this, the order parameters were not significantly different than 0 in any case. This suggests that the cells on substrates aligned with cells that were near to them even if they were not in direct contact. Interestingly, this was not seen for cells in hanging drops, suggesting longer-range cell alignment occurs only in monolayer clusters on solid substrates.

## DISCUSSION

Understanding actomyosin contractility is one key aspect of cell mechanics that is required for explaining the dynamics of tissue remodeling. Direct experimental measurements of force generation in monolayer clusters without a rigid surface can aid in our understanding of the remodeling of epithelial monolayers. The objective of this study was to develop a new experimental methodology and analysis framework in order to provide an approach to understanding the mechanodynamics of unsupported epithelial monolayers. The lack of a solid support provided a less complicated system that allowed for remodeling not possible on a solid substrate. This method also provides a rational approach to teasing apart substrate and intercellular connections.

As a validation of the practicality of the hanging drop model, we investigated the effect of cytoskeletal drugs on the deformation of MDCK monolayer clusters in both the absence and presence of mechanically supporting substrates. By comparing cell response in these two support regimes, we were able to identify the intrinsic contractile dynamics of an epithelial monolayer cluster. Since contraction/dilation was muted on solid supports, removing the solid substrate allowed the monolayer to clusters contract/dilate. The deformation we observed by tracking nuclei was echoed in the deformation of the individual cell and the monolayer cluster outlines. This verified the use of nuclei as an effective cluster deformation marker. The observed deformations established that the contraction and dilation of monolayer clusters could be directly observed in the absence of a substrate by tracking cell nuclei in hanging drop clusters.

To calculate stress in our droplets we used the mean strains obtained from our data and the Young’s modulus reported by Harris et al. for MDCK monolayers of 20±2kPa (Harris *et al*, 2012). This results in an apparent decrease of stress of 320±50Pa for the blebbistatin treatment and an apparent increase of stress of 190±40Pa for the CalA treatment. There was a negligible change in stress for the control case. In other experiments, MDCK cells grown on pillars have been found to have average traction stresses of around 800Pa in a monolayer (Roure *et al*, 2005). Our stresses are smaller than, but of the same order of magnitude as, stresses observed in cells grown on pillars. The calculated stresses are reasonable considering that the hanging drop is expected to support less traction force than a stiffer substrate. This demonstrates that the hanging drop system can be used to calculate the intrinsic stress produced by actomyosin contraction and relaxation in monolayer clusters.

Furthermore, varying the cluster substrate allowed us to relate contractile dynamics to substrate rigidity. Our hanging drop method allowed us to measure the intrinsic remodeling that can occur in response to actomyosin dynamics. This remodeling was muted by the presence of a solid substrate, demonstrating that cluster contraction is significantly restrained when bound to a rigid surface. In general, stiff substrates increase cell-substratum adhesion (Pelham, 1998; Engler *et al*, 2003; Yeung *et al*, 2005; Ghibaudo *et al*, 2008) and increase traction forces (Maruthamuthu, 2011). It has also been shown that compliant substrates promote focal adhesions that are dynamic and irregular in shape while stiff substrates promote the formation of stable arrays of elongated focal adhesions (Pelham, 1998). When the cells are grown on glass, a large portion of the work from the actomyosin contraction was converted into substrate deformation instead of cellular remodeling. On softer substrates (PA gels in particular) cells are still restrained, however the smaller, more dynamic focal adhesions may allow for more movement. In this case, a smaller portion of the work will go into deforming the substrate. The contraction energy of the hanging drop cells is expected to be smaller than the contraction energy of cells on glass or PA. However, the cells can move more freely in the cluster because they are not attached by focal adhesions to a substrate that is difficult to deform. Only a negligible amount of work should go into deforming the basement membrane-like sheet below the monolayer cluster, allowing for the larger cellular remodeling events observed in this paper.

Interestingly, when the clusters were grown on PA gels there was a correlation between major strain direction and nuclear alignment. However, for cells grown on at the air-water interface in a hanging drop, the direction of the major strain was independent of nuclear alignment. The nuclei of single epithelial cells grown on flat rigid surfaces have been previously found to align with the actin cytoskeleton and the major length axis of the cell (McKee *et al.*, 2011; Raghunathan *et al.*, 2013). This suggests that the actin fibers are aligned with the major strain direction when on PA gels. This is congruous with the theory that individual cells are contracting and relaxing preferentially in the direction of their actin cytoskeletons resulting in a global deformation along the cluster axis that many cells were aligned with. It has also been shown that matrix stiffness aids in regulating the polarization and alignment of stress fibers within cells (Zemel *et al.*, 2010). The lack of correlation between nuclear orientation and strain in the hanging drop may be due to reduced alignment of actin within each cell and the reduction in longer-range correlated cell-cell alignment that was observed on rigid substrates. In the case of the stiff glass substrate, although there should be greater alignment due to the substrate, focal adhesions pinned cells to the substrate leading to null cluster strains, making it arduous to correlate to strain. The softer PA substrate allowed for substrate mediated alignment and limited cell movement permitting a correlation between cluster strain and nuclear alignment to be observed.

Remarkably, a solid substrate appears to be required for longer-range cell-cell alignment. By calculating nuclear order parameters, we found that cells in hanging drops were only aligned with their nearest neighbors, while cells on substrates were aligned with their next nearest and third nearest neighbors as well. This longer-range organization may be a result of substrate cues not present in hanging drops. It suggests also that signalling is necessary to develop long-range stresses. One possibility is that long-range orientational correlations are the result of elastic coupling of cells mediated by deformation of the compliant substrate. This is consistent with work showing that intercellular forces develop within the cluster of cells grown on a gel (Mertz *et al.*, 2013). This unexpected finding was made evident by comparison with the extreme condition of an air-water interface substrate. It would not have been possible without the novel hanging drop monolayer. Our new methodology has revealed some surprising and novel insights into the mechanodynamics of epithelial monolayers. However, it also raises several new questions that will warrant further study in the future.

For instance, do the cells in the monolayer establish apico-basal polarization? Is this polarization triggered without a solid substrate? This could be determined with more advanced staining techniques. Are the same structures formed on planar non-adhesive substrates or is the curved drop shape needed? Plating cells in dishes with non-adhesive coatings could be a method to investigate this. How do clusters grown on substrates recoil when detached? This could be investigated using trypsin to detach the clusters from the substrate or by using thermo-sensitive polyacrylamide derivatives to allow release of the sheets. Future work will include computational modeling of these scenarios to examine these effects in further detail. These hanging drop monolayers can also be made with epithelial cells stably expressing a fluorescently marked protein of interest and could also be utilized for a broad range of mechanical experiments investigating substrate effects on monolayers. Novel tools and methodologies for understanding the mechanics of monolayers will enable the advancement of the biophysics field.

## MATERIALS AND METHODS

### Cell culture

Madin-Darby Canine Kidney (MDCK) epithelial cells were cultured in DMEM with 10% fetal bovine serum (FBS), 50mg/ml streptomycin and 50U/ml penicillin antibiotics (all from Hyclone Laboratories Inc.). Cells were cultured at 37μC in a 5% CO_2_ incubator on 100mm tissue culture dishes (Corning). In some experiments, cells were cultured on solid substrates. In these cases, 1x10^5^ cells were cultured in 34mm tissue culture dishes (TPP). The dishes either contained a flexible (soft) polyacrylamide (PA) hydrogel coating, previously described (Wang *et al.*, 2000; Kandow *et al.*, 2007; Tse and Engler, 2010), or a bare glass coverslip bottom. To produce a PA gel with a young’s modulus of 4.8kPa (Quinlan and Billiar, 2012), 7.5% acrylamide was polymerized with 0.053% bisacrylamide on (3-Aminopropyl)trimethoxysilane coated coverslips with 1mg/ml ammonium persulfate (Biorad) and 0.15% tetramethylethylenediamine (Biorad). Prior to cell seeding, the hydrogel surface was functionalized with 100*μ*g/mL collagen (Gibco) in PBS using 0.5mg/mL sulfo-sanpah (Pierce) as a crosslinker. In some experiments, cells were cultured in hanging drops. MDCK cells were trypsinized and then resuspended in media at a low dilution of 4000 cells/ml. The cells were then placed in hanging drops of 40*μ*l on the lower surface of the lids of plastic Petri dishes containing Phosphate Buffered Saline (PBS). Hanging drops were imaged after 24h of culture.

### Drug treatments

Cells were treated with blebbistatin (Sigma) dissolved in DMSO at a final concentration of 40μM or calyculin A (CalA, Sigma) dissolved in water at a final concentration of 300nM for 30 minutes in an incubator at 37*μ*C and 5% CO_2_ between before and after images.

### Imaging

All images were acquired on a TiE A1-R laser scanning confocal microscope (LSCM) (Nikon) with a 40X long working distance objective. Images were acquired with a standard LSCM configuration with appropriate laser lines and filter blocks. Cell nuclei were stained with Hoechst. In some cases, cells expressing GFP-actin were produced with BacMam reagent (Invotrogen, CellLight^®^ Reagent BacMam 2.0) using manufacture protocols at 20 particles per cell. Cells were plated and incubated for at least 48h before being placed into hanging drops using the protocol above. Cells were also stained with wheat germ agglutinin (WGA) coupled to Texas Red-X (Invitrogen) to reveal the cell membrane. Cells stained on glass were fixed with 3.5% paraformaldehyde and permeabilized with Triton X-100 at 37°C. Actin was stained with Phalloidin Alexa Fluor 546 (Invitrogen). Following this vinculin was stained with monoclonal anti-vinculin primary antibody (Sigma) and Alexa Fluor 488 rabbit anti-mouse immunoglobin (Invitrogen) secondary antibody. DNA was labeled with DAPI (Invitrogen).

### Strain Quantification

The strain in the monolayer was determined with the displacements of the cell nuclei during a 30 minute treatment. This method was validated by tracking the cell membrane boundary of the cluster and by tracking actin in the cell margins. Images were processed and analyzed with ImageJ. Maximum intensity z-projections of 10, 5μm thick slices were used for strain calculations with no enhancements applied (Figure 2A). The positions of the nuclei in these images were determined using the analyze particles function after thresholding. The positions of the nuclei in the before and after images were manually linked and analyzed using Matlab. A Matlab program was used to account for differences in dish placement in the before and after images, as well as to determine nuclear displacements and calculate the strain tensor of the cluster.

In this program, nuclei positions were translated in both images so that the center of the nuclei was at the origin. These translated positions were then rotated so that the sum of the nuclei displacements were minimized (Fig 1B). This process was needed so that there were no artificially large displacements from the movement of the dish to and from an incubator between images. The initial positions (x, y) and displacements (u, v) of the nuclei were then used to fit the cluster with least squares to the displacement equations: u=c_1_x+c_2_y+c_3_xy+c_4_x^2^+c_5_y^2^ and v=c_6_x+c_7_y+c_8_xy+c_9_x^2^+c_10_y^2^ (Bathe, 2006). Second order terms were included to account for non-constant strain throughout the cluster. These equations were then used to determine the diagonal components of the Green-Lagrangian strain tensor,

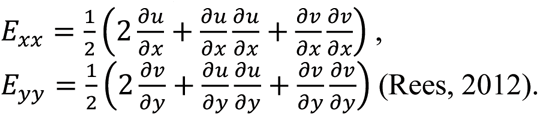

Substitution of our displacement equations resulted in

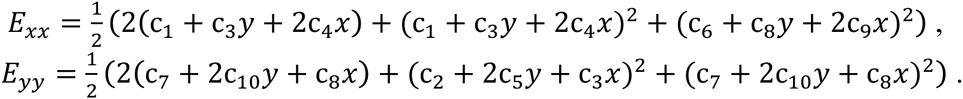

These were used along with the shape of the cluster to determine the average x-axis extensional strain, *E*_*xx*_, and average y-axis extensional strain, *E*_*yy*_, in the cluster. The cluster was then rotated to find the principle axes by maximizing the difference between the y-axis strain and the x-axis strain. After this rotation, the y-axis strain was the major principle strain (E_1_) and the x-axis strain was the minor principle strain (E_2_) (Fig 1C). The mean of the principle strains, 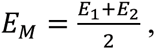, is the deformation due to a change in area (Mase *et al.*, 2009).

For experiments measuring the deformation of the cluster outline, points along the cluster outline labeled in the cell membrane stained images were tracked and used in the code above in place of nuclei positions. In some experiments cellular-level strains were determined in cells expressing actin-GFP. Cell deformation was determined by fitting cell boundaries with polygons. The displacements of the vertices were least squares fit to the displacement equations u=c_1_x+c_2_y and v=c_3_x+c_4_y to impose uniform strain inside the cell (Bathe, 2006). Only first order terms in the displacements u and v were required for these calculations because the individual cells were much smaller than the cluster. The strain in each cell was calculated using the principle strain axes of the cluster determined using the nuclei displacements.

### Order Parameter Calculations

Orientational order parameters were calculated using the angles between different cluster features. The nematic order parameter was used, which in two dimensions is *S*= 〈2 cos^2^ *θ* — 1〉 (Mercurieva and Birshtein, 1992). The order parameter will be 1 if the features are completely aligned, −1 if they are completely antialigned, and 0 if there is no alignment. The nuclei in the thresholded before images were fitted to ellipses in Matlab to determine the orientations of the major length axes and the eccentricities of the nuclei. The orientation of the major length axis of the cluster was determined using the positions of the nuclei and finding the axis that minimized the square of the distances between the axis and the nuclei. Four different types of order parameters were calculated. Nuclei-strain order parameters (*S*_*ns*_) were determined using the angles between the major length axes of the nuclei and the major strain axis of the cluster. Nuclei-cluster orientation order parameters (*S*_*nl*_) were calculated using the angles between the major length axes of the nuclei and the cluster major length axis. Cluster orientation-strain order parameters (*S*_*ls*_) were also calculated with the angles between the major length and major strain axes of clusters. In addition, nuclei-nuclei order parameters (*S*_*nn*_) were calculated using the angles formed between the major axis of each nucleus and the major axis of its nearest neighbor. This was also repeated for next nearest neighbors, third nearest neighbors and fourth nearest neighbors.

## ACKNOWLEDGMENTS

This work was supported by a Natural Sciences and Engineering Research Council (NSERC) Discovery Grant. C.G. was supported by the uOttawa NSERC-CREATE program in Quantitative Biomedicine. A.E.P. gratefully acknowledges generous support from the Canada Research Chairs (CRC) program.

## REFERENCES

Bartosh, T. (2010). Aggregation of human mesenchymal stromal cells (MSCs) into 3D spheroids enhances their antiinflammatory properties. Proc. Natl. Acad. Sci. USA 107, 1–6.

Bathe, K.-J. (2006). Finite Element Procedures.

Chu, Y.-S., Thomas, W. a, Eder, O., Pincet, F., Perez, E., Thiery, J. P., and Dufour, S. (2004). Force measurements in E-cadherin-mediated cell doublets reveal rapid adhesion strengthened by actin cytoskeleton remodeling through Rac and Cdc42. J. Cell Biol. 167, 1183–1194.

Discher, D. E., Janmey, P., and Wang, Y.-L. (2005). Tissue cells feel and respond to the stiffness of their substrate. Science 310, 1139–1143.

Eckhart, L., Reinisch, C., Inoue, S., Messner, P., Dockal, M., Mayer, C., and Tschachler, E. (2003). A basement membrane-like matrix formed by cell-released proteins at the medium/air interface supports growth of keratinocytes. Eur. J. Cell Biol. 82, 549–555.

Engler, a., Sheehan, M., Sweeney, H. L., and Discher, D. E. (2003). Substrate compliance vs ligand density in cell on gel responses. Eur. Cells Mater. 6, 7–8.

Fernandez-Gonzalez, R., Simoes, S. D. M., Röper, J.-C., Eaton, S., and Zallen, J. a (2009). Myosin II dynamics are regulated by tension in intercalating cells. Dev. Cell 17, 736–743.

Ghibaudo, M., Saez, A., Trichet, L., Xayaphoummine, A., Browaeys, J., Silberzan, P., Buguin, A., and Ladoux, B. (2008). Traction forces and rigidity sensing regulate cell functions. Soft Matter 4, 1836.

Ghibaudo, M., Trichet, L., Le Digabel, J., Richert, A., Hersen, P., and Ladoux, B. (2009). Substrate topography induces a crossover from 2D to 3D behavior in fibroblast migration. Biophys. J. 97, 357–368.

Harris, A. R., Bellis, J., Khalilgharibi, N., Wyatt, T., Baum, B., Kabla, A. J., and Charras, G. T. (2013). Generating suspended cell monolayers for mechanobiological studies. Nat. Protoc. 8, 2516–2530.

Harris, A. R., Peter, L., Bellis, J., Baum, B., Kabla, A. J., and Charras, G. T. (2012). Characterizing the mechanics of cultured cell monolayers. Proc. Natl. Acad. Sci. USA 109, 16449–16454.

Heisenberg, C.-P., and Bellaïche, Y. (2013). Forces in tissue morphogenesis and patterning. Cell 153, 948–962.

Huh, D., Matthews, B. D., Mammoto, A., Montoya-Zavala, M., Hsin, H. Y., and Ingber, D. E. (2010). Reconstituting organ-level lung functions on a chip. Science 328, 1662– 1668.

Inoue, S., Reinisch, C., Tschachler, E., and Eckhart, L. (2005). Ultrastructural characterization of an artificial basement membrane produced by cultured keratinocytes. J. Biomed. Mater. Res. A 73, 158–164.

Kandow, C. E., Georges, P. C., Janmey, P. a, and Beningo, K. a (2007). Polyacrylamide hydrogels for cell mechanics: steps toward optimization and alternative uses. Methods Cell Biol. 83, 29–46.

Last, J., Liliensiek, S., Nealey, P., and Murphy, C. (2009). Determining the mechanical properties of human corneal basement membranes with atomic force microscopy. J. Struct. Biol. 167, 19–24.

Lecuit, T., and Lenne, P.-F. (2007). Cell surface mechanics and the control of cell shape, tissue patterns and morphogenesis. Nat. Rev. Mol. Cell Biol. 8, 633–644.

Lemmon, C. a, Chen, C. S., and Romer, L. H. (2009). Cell traction forces direct fibronectin matrix assembly. Biophys. J. 96, 729–738.

Leptin, M. (2005). Gastrulation movements: The logic and the nuts and bolts. Dev. Cell 8, 305–320.

Martin, A. C., Gelbart, M., Fernandez-Gonzalez, R., Kaschube, M., and Wieschaus, E. F. (2010). Integration of contractile forces during tissue invagination. J. Cell Biol. 188, 735– 749.

Maruthamuthu, V. (2011). Cell-ECM traction force modulates endogenous tension at cell–cell contacts. Proc. Natl. Acad. Sci. USA.

Mase, G. T. E., Smelser, R. E., and Mase, G. T. E. (2009). Continuum Mechanics for Engineers, Third Edition.

McKee, C. T., Raghunathan, V. K., Nealey, P. F., Russell, P., and Murphy, C. J. (2011). Topographic modulation of the orientation and shape of cell nuclei and their influence on the measured elastic modulus of epithelial cells. Biophys. J. 101, 2139–2146.

Mercurieva, A. a., and Birshtein, T. M. (1992). Liquid-crystalline ordering in two-dimensional systems with discrete symmetry. Die Makromol. Chemie, Theory Simulations 1, 205–214.

Mertz, A. F., Che, Y., Banerjee, S., Goldstein, J. M., Rosowski, K. a, and Revilla, S. F. (2013). Cadherin-based intercellular adhesions organize epithelial cell–matrix traction forces. Proc. Natl. Acad. Sci. 110, 842–847.

Nelson, C. M., Jean, R. P., Tan, J. L., Liu, W. F., Sniadecki, N. J., Spector, A. a, and Chen, C. S. (2005). Emergent patterns of growth controlled by multicellular form and mechanics. Proc. Natl. Acad. Sci. USA 102, 11594–11599.

Ng, M. R., Besser, A., Brugge, J. S., and Danuser, G. (2014). Mapping the dynamics of force transduction at cell-cell junctions of epithelial clusters. Elife 4, e03282.

Notbohm, J., Kim, J. H., Asthagiri, a. R., and Ravichandran, G. (2012). Three-dimensional analysis of the effect of epidermal growth factor on cell-cell adhesion in epithelial cell clusters. Biophys. J. 102, 1323–1330.

Owaribe, K., Kodama, R., and Eguchi, G. (1981). Demonstration of contractility of circumferential actin bundles and its morphogenetic significance in pigmented epithelium in vitro and in vivo. J. Cell Biol. 90, 507–514.

Pelham, R. J. (1998). Cell Locomotion and Focal Adhesions Are Regulated by the Mechanical Properties of the Substrate. Biol. Bull., 348–350.

Pelham, R., and Wang, Y. (1997). Cell locomotion and focal adhesions are regulated by substrate flexibility. Proc. Natl. Acad. Sci. USA 95.

Quinlan, A. M. T., and Billiar, K. L. (2012). Investigating the role of substrate stiffness in the persistence of valvular interstitial cell activation. J. Biomed. Mater. Res. A 100, 2474–2482.

Raghunathan, V. K., McKee, C. T., Tocce, E. J., Nealey, P. F., Russell, P., and Murphy, C. J. (2013). Nuclear and cellular alignment of primary corneal epithelial cells on topography. J. Biomed. Mater. Res. - Part A 101 A, 1069–1079.

Rauzi, M., Lenne, P.-F., and Lecuit, T. (2010). Planar polarized actomyosin contractile flows control epithelial junction remodelling. Nature 468, 1110–1114.

Rauzi, M., Verant, P., Lecuit, T., and Lenne, P.-F. (2008). Nature and anisotropy of cortical forces orienting Drosophila tissue morphogenesis. Nat. Cell Biol. 10, 1401–1410.

Rees, D. (2012). Basic Engineering Plasticity: An Introduction with Engineering and Manufacturing Applications.

Roh-Johnson, M., and Shemer, G. (2012). Triggering a Cell Shape Change by Exploiting Pre-Existing Actomyosin Contractions. Science (80-.). 335, 1232–1235.

Roure, O. Du, Saez, A., Buguin, A., Austin, R. H., Chavrier, P., Silberzan, P., and Ladoux, B. (2005). Force mapping in epithelial cell migration. Proc. Natl. Acad. Sci. USA 102, 2390–2395.

Solon, J., Levental, I., Sengupta, K., Georges, P. C., and Janmey, P. a (2007). Fibroblast adaptation and stiffness matching to soft elastic substrates. Biophys. J. 93, 4453–4461.

Tchao, R. (1989). Epithelial cell interaction in air-liquid interface culture. Vitr. Cell. Dev. Biol. 25.

Tremblay, D., Andrzejewski, L., Leclerc, A., and Pelling, A. E.(2013). Actin and microtubules play distinct roles in governing the anisotropic deformation of cell nuclei in response to substrate strain. Cytoskeleton 70, 837–848.

Trichet, L., Le Digabel, J., Hawkins, R. J., Vedula, S. R. K., Gupta, M., Ribrault, C., Hersen, P., Voituriez, R., and Ladoux, B. (2012). Evidence of a large-scale mechanosensing mechanism for cellular adaptation to substrate stiffness. Proc. Natl. Acad. Sci. USA 109, 6933–6938.

Tse, J. R., and Engler, A. J. (2010). Preparation of hydrogel substrates with tunable mechanical properties. Curr. Protoc. Cell Biol. 10, 16.

Vicente-Manzanares, M., Ma, X., Adelstein, R. S., and Horwitz, A. R. (2009). Non-muscle myosin II takes centre stage in cell adhesion and migration. Nat. Rev. Mol. Cell Biol. 10, 778–790.

Vogel, V., and Sheetz, M. (2006). Local force and geometry sensing regulate cell functions. Nat. Rev. Mol. Cell Biol. 7, 265–275.

Wang, Y. N., Galiotis, C., and Bader, D. L. (2000). Determination of molecular changes in soft tissues under strain using laser Raman microscopy. J. Biomech. 33, 483–486.

Wiebe, C., Wayne, B. G., and Brodland, G. W. (2005). Tensile properties of embryonic epithelia measured using a novel instrument. J. Biomech. 38, 2087–2094.

Yeung, T., Georges, P. C., Flanagan, L. a., Marg, B., Ortiz, M., Funaki, M., Zahir, N., Ming, W., Weaver, V., and Janmey, P. a. (2005). Effects of substrate stiffness on cell morphology, cytoskeletal structure, and adhesion. Cell Motil. Cytoskeleton 60, 24–34.

Yonemura, S., Itoh, M., Nagafuchi, a, and Tsukita, S. (1995). Cell-to-cell adherens junction formation and actin filament organization: similarities and differences between non-polarized fibroblasts and polarized epithelial cells. J. Cell Sci. 108 ( Pt 1, 127–142.

Zegers, M. M. P., O’Brien, L. E., Yu, W., Datta, A., and Mostov, K. E. (2003). Epithelial polarity and tubulogenesis in vitro. Trends Cell Biol. 13, 169–176.

Zemel, a., Rehfeldt, F., Brown, a. E. X., Discher, D. E., and Safran, S. a. (2010). Optimal matrix rigidity for stress-fibre polarization in stem cells. Nat. Phys. 6, 468–473.

